# RESUPER: a new computer program for identifying ordered regions, superposition ranges and conformational states in protein ensembles

**DOI:** 10.64898/2026.11.22.768412

**Authors:** Yukthika Boddeda, Woonghee Lee

## Abstract

NMR structures are deposited as ensembles of conformers, and their precision is reported after superposition on a chosen set of residues, a choice that is difficult for proteins with several domains or flexible regions. We present RESUPER, a program that identifies rigid domains, selects a superposition range for each domain, and detects distinct conformational states in protein ensembles. RESUPER follows the two-stage method of CYRANGE, but each domain split is decided by a geometric test that can be checked on the structure. On two test sets fixed before evaluation, RESUPER counted the domains correctly in 95 of 96 and 45 of 48 ensembles (CYRANGE: 76 and 39) and put two separate rigid bodies into one range in 0 and 4 ensembles (CYRANGE: 17 and 9). On 222 randomly selected NMR entries, the two programs gave the same number of domains for 88% of the entries, and none of four prespecified comparisons, applied post hoc to CYRANGE, showed a statistically significant difference. For apo-Pin1 (PDB 7SA5), RESUPER found the two deposited conformational states. Analyzed separately, one state has a single superposition range covering both domains and the other has separate ranges. RESUPER is integrated into POKY with a graphical user interface.

## Introduction

Structure ensembles from NMR spectroscopy, molecular dynamics or ensemble refinement are compared after superposition, and the residues used for the superposition determine what the RMSD measures. If the superposition includes a mobile terminus, the RMSD mostly reflects that terminus. If it is restricted to a rigid part of the protein, the RMSD describes how well that part is defined. Choosing the residues automatically requires finding the rigid domains, measuring the mobility of each residue, and selecting the range over which the fit is computed.

Kirchner and Güntert^1^ showed that this choice can be made objectively with a two-stage method, implemented in CYRANGE. Rigid domains are first identified from torsion-angle order parameters and the variances of interatomic distances, and the residue range of each domain is then refined. CYRANGE is the established tool for this task, and the wwPDB runs it on every NMR entry^2^. The first stage leaves one decision open: where to stop dividing the protein. A hierarchical clustering tree of the well-ordered atoms offers a candidate partition at every level. The published objective ranks these levels but gives no absolute test for accepting one, so where the division stops depends on an empirically chosen constant. Errors at this step are hard to see. A range that spans two rigid bodies moving relative to each other still gives a plausible RMSD, so the wrong decomposition cannot be recognized from the reported number. Nor does any rule prevent a split of a region that is simply flexible. A small hinge can also stay hidden inside a range that looks ordered until exactly the right residues are removed; removing one residue too many breaks up the domain.

Residue ranges also do not tell whether an ensemble is a single population with thermal motion or a mixture of discrete conformational states^3,4^. A multi-state ensemble can have well-defined rigid domains in every state, so even a correct superposition range says nothing about the number of states. The conformers of different states are often interleaved in model order, so the states do not appear as a block of consecutive models either.

CYRANGE is distributed as a closed-source binary for a limited set of x86-64 Linux platforms. It cannot run on ARM64 workstations and clusters, which are now common, without emulation. Its source code is not available, so users cannot inspect or extend it. RESUPER keeps the two-stage framework but replaces each tuned step of domain selection with a test that has a geometric meaning, so every domain boundary can be checked against a quantity measured on the structure. It counts conformational states from the same distance information, and it is a portable Python program. RESUPER is included in POKY^5^, which provides a graphical interface, writes the selected ranges in Xplor-NIH, CYANA and PyMOL formats, and can save each detected state as a separate superposed ensemble (Figure 1; two-letter code *yS*).

**Figure 1.**
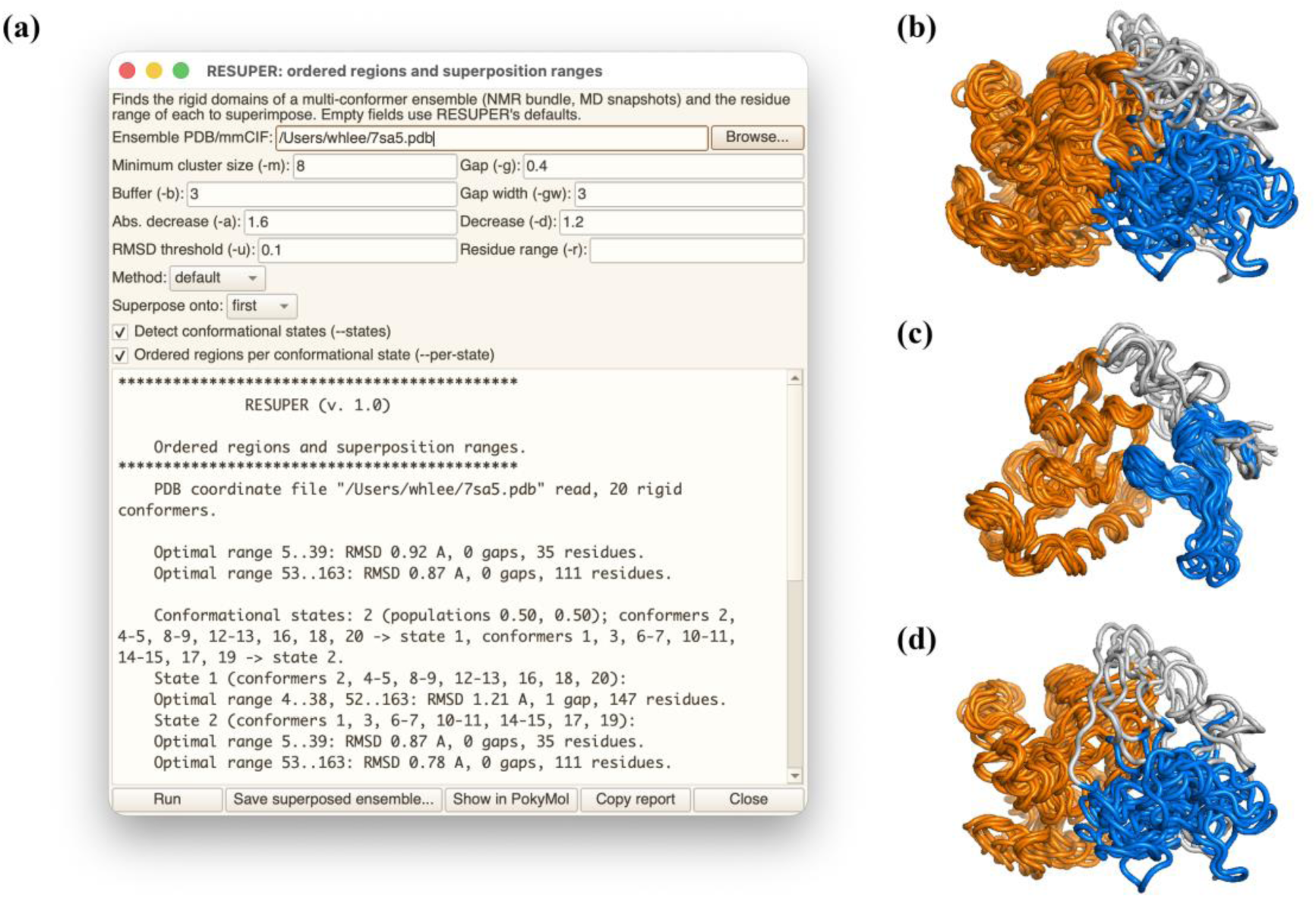
RESUPER in POKY (two-letter code *yS*), applied to apo-Pin1 (PDB 7SA5). (a) The RESUPER dialog with the CYRANGE parameters, the options “Detect conformational states” and “Ordered regions per conformational state”, and the report: the two ranges of the whole ensemble, the two states with their conformers given as the model numbers of the input file, and the ranges of each state. (b) All 20 conformers. (c) State 1 (10 conformers). (d) State 2 (10 conformers). The PPIase domain is shown in orange, the WW domain in blue, and the residues outside the reported ranges in white.

Within POKY^6^, RESUPER will be applied broadly as part of the integrative NMR platform^7^. In the automated structure calculation of PONDEROSA-C/S^8^, with AUDANA^9^ for solution NMR data and AUDASA^10^ for solid-state NMR data, it will check the ordered regions of the calculated ensemble at every cycle of the calculation.

## Computational Methods

### Overview

RESUPER follows the two-stage method of Kirchner and Güntert^1^: rigid domains are identified first, and the superposition range of each domain is then refined. The default parameters are listed in Table 1.

**Table 1.** Default parameters of RESUPER.

| Symbol | Meaning | Default |
|---|---|---|
| τ | Minimum split score | 0.12 Å |
| *g* | Merge-height ratio, block test | 7 |
| *d* | RMS motion, block test | 0.5 Å |
| μ | Minimum domain size | 8 core atoms |
| *m* | Start-range extension | 3 residues |
| γ | Gap-opening weight | 0.4 |
| δ<sub>abs</sub> | Absolute decrease gate | 1.6 Å |
| δ | Relative decrease base | 1.2 |
| *q̂*<sub>2</sub> | Silhouette threshold for more than one state | 0.60 |

### Core atoms and clustering

For a dihedral angle θ in *N* conformers, the angular order parameter is

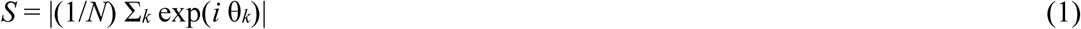

*S* is 1 when the angle is the same in all conformers and approaches 0 when the angle is randomly distributed. All rotatable dihedrals except ω are used. A residue with at least one torsion with *S* ≥ *S*_cut_ is a core residue, and its Cα is a core atom. *S*_cut_ is the value *S_i_* that maximizes^1^

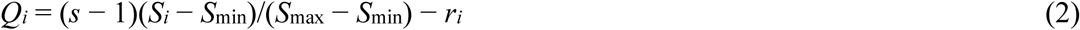

where *s* is the number of torsions and *r_i_* is the rank of *S_i_*. The core atoms are clustered by Ward linkage^11^ on the distance variances *V_ij_* = σ(*d_ij_*)^2^ over the conformers, so atoms whose mutual distances change little are grouped together.

### Domain split criterion

Each node of the clustering tree has two children, *L* and *R*. Let *r*(·) denote the backbone (N, Cα, C′) RMSD to the mean structure of a group of atoms superimposed on its own. The split score is

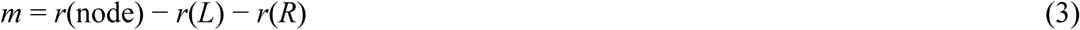

*m* is not itself the decrease in RMSD obtained by fitting the two parts separately. The pooled RMSD of the separately fitted parts never exceeds *r*(*L*) + *r*(*R*), so *m* is never larger than that decrease, and the test is conservative.

A domain must contain at least μ = 8 core atoms, and *C* is the total number of core atoms. (i) If both children have at least μ atoms, the node is split when *m* ≥ τ (0.12 Å) or when *V* shows two blocks in relative motion: the merge height of the node is at least 7 times those of its children, and the RMS motion between the two parts is at least 0.5 Å. (ii) A child with fewer than μ atoms is treated as noise and discarded. (iii) If both children have fewer than μ atoms, the node is a domain only if it is rigid; it is discarded as flexible when *r*(node) − max[*r*(*L*), *r*(*R*)] ≥ τ. (iv) A node whose own split fails is still divided if a split that passes the test, with parts of at least ⌈*C*/8⌉ core atoms, exists further down the tree. The groups that remain are the domains.

### Start ranges and refinement

Each domain starts from the span of its core residues, extended by three residues on each side. The range is then refined in cycles. Isolated residues are removed, and the residue with the largest average displacement is removed if the decrease in RMSD, Δ*r*, weighted for any gap the removal opens, satisfies Δ*r* ≥ 1.6 Å·*n*/*N* or Δ*r*/*r* ≥ (1.2 + 3.0/*M*)·*n*/*N*, where *n*/*N* is the fraction of atoms removed and *M* is the number of selected residues. The cycles stop when the range no longer changes, and gaps shorter than three residues are then filled. All RMSDs use the N, Cα and C′ atoms after SVD superposition^12^.

### Multiconformer detection

To estimate the number of conformational states^3,4^, the Cα–Cα distance matrix of each conformer is reduced to three principal components. The conformers are clustered by Ward linkage in this space, and the tree is cut into *k* = 2, 3, … groups, up to the largest *k* at which every group still has at least three conformers. Each partition is scored by its silhouette coefficient^13^, which measures how well the groups are separated; a single group is given a score of 0. The number of states is the value of *k* with the largest increase in silhouette over *k* − 1. It is reported only if the silhouette of that partition is at least 0.60; otherwise the ensemble is reported as one state. The 0.60 threshold was calibrated on four Pin1 ensembles and single-state controls.

Conformers are identified by their model numbers in the input file. When conformers of different states are interleaved in model order, each state is printed as a list of runs (e.g., 1, 3, 6–7). With `--per-state`, the domain identification and range refinement are repeated on the conformers of each state, and the ranges of each state are reported below those of the whole ensemble.

### Implementation

RESUPER is a Python package with a command-line interface that uses CYRANGE’s parameter names and units `(-a, -b, -d, -g, -gw, -m, -r, -l, -u`); the repository version of 22 November 2026 was used here. `--states` adds the state report, `--per-state` adds the ranges of each state, `--write-ensemble` writes the superposed ensemble, and `--split-states` writes one file per state. PDB and mmCIF input are accepted. `--method reference` is an approximate reimplementation of CYRANGE 2.0, built from the published description^1^ and adjusted against reports of the distributed program on the development ensembles. It is used only for comparison. Its results are not those of CYRANGE, and every comparison with CYRANGE in this work uses the distributed program or its results in the wwPDB. Defective input is reported instead of being repaired silently. HETATM records and non-standard residues are excluded from the protein, and chain breaks keep ranges discontinuous. Missing atoms, non-numeric coordinates, malformed lines, unnumbered MODEL records, single-model input and input with no usable atoms give either a result or an explicit error (`tests/test_robust.py`). In POKY^5^, the same command-line program produces the report, the range strings for Xplor-NIH, CYANA and PyMOL, and the per-state ensembles (Figure 1).

### Benchmarks with known domains

RESUPER was compared with CYRANGE 2.0 and with the reference mode on the following ensembles. The first set was 13 synthetic ubiquitin ensembles. The second was 134 real NMR ensembles collected during development, each with a report from the distributed CYRANGE 2.0 binary: curated single- and multi-domain cases, a frozen list of 99 entries selected from the PACSY database^14,15^, and two-state bundles built from the AISAR ensembles^4^. These 134 ensembles are an evaluation set, not a random sample. Two held-out development sets of 96 ensembles each had hinge and disorder motions applied in torsion space. Two test sets were run only after the method was frozen, with success criteria fixed in advance: set 3 (96 ensembles) and set R (48 ensembles made by applying motions to twelve real ensembles). Perturbed copies of the 134 ensembles had reversed model order, two subsets of 80% of the models, or 0.10 Å and 0.25 Å of coordinate noise. Finally, 788 ensembles with one to four bodies were built on 40 proteins; 160 of them contain two or more bodies. The measures were the number of domains, the Jaccard index (the overlap between a predicted and a true domain), coverage, contamination by flexible residues, and mixed domains (domains with ≥ 3 residues and ≥ 10% from a second true domain).

### Comparison with CYRANGE results in the wwPDB

The wwPDB runs CYRANGE on every NMR entry and stores the domain of each residue in the validation report^2^. Domains were read from the first model of each report. For all 108 repository entries that have both a saved CYRANGE 2.0 report and a PDB ID, these domains are identical to those of the distributed binary (9/9 of the multi-domain entries), so both comparisons use the same reference. Entries with solution NMR structures, ≥ 10 models and one protein chain, and not used in RESUPER development, were taken from the RCSB PDB^16^. Before any program was run, they were grouped with CATH^17^, SCOP^18^ and ECOD^19^ annotations: two (A) or ≥ 3 (B) annotated domains; single-domain controls with at least 20 residues and 20% of the chain outside the annotated domain (C) or with an 8–25-residue helix outside it (D); and helical peptides of 25–50 residues (E). Two random samples were drawn with fixed seeds before any download (60 entries, seed 20261111; 166 entries, seed 20261113) and screened by fixed rules. Four of the 226 entries were removed because their conformers differ in backbone atoms or their chains are largely non-standard (2K60, 2OFG, 9LYH, 9XFN), leaving 60 and 162 entries. Every tested entry had a CYRANGE result in its wwPDB report.

Domains of the two programs were paired by Jaccard index with the Hungarian algorithm^20^. An annotated domain was recovered when ≥ 50% of it lay in one unmixed domain and missed when < 20% of it was covered. A domain found by only one program passed the rigid-domain check if its RMSD was ≤ 1.0 Å and its rotation relative to the rest of the protein exceeded the 95th percentile of a size-matched baseline. Four hypotheses were tested: recovered annotated domains (H1), missed annotated domains (H2), over-splitting of controls (H3), and short helices reported as domains (H4). The tests were Wilcoxon^21^ and exact McNemar^22^ tests, two-sided at α = 0.05, as specified in advance in the decision log. No correction for the four comparisons was specified, and none of the p values is close to the threshold. The hypotheses were specified with the reference mode as the comparator, so their use with CYRANGE is post hoc. Within the five groups, this random-sample comparison is the estimate least affected by selection bias in this study. The illustrative cases below were selected for a property and do not show how often it occurs.

### Selection of illustrative cases

All remaining entries of the pool were compared with CYRANGE. Entries where the programs differed were judged by rules fixed before the scan: annotation, rigidity of the CYRANGE domains, and whether RESUPER-only domains were rigid and moving. An entry was excluded if the same evidence favored CYRANGE. Two categories were then defined. In category A, RESUPER reports more domains than CYRANGE, all of ≥ 20 residues and rigid (≤ 1.0 Å), with no split inside one helix and no contradiction in the original publication. In category B, every RESUPER domain matches a CYRANGE domain (Jaccard ≥ 0.9), the remaining CYRANGE domains are not rigid (> 1.0 Å, ≥ 8 residues), the protein has ≥ 60 residues, and the original publication explicitly describes that region as disordered or flexible. These categories show cases in which the evidence favors RESUPER; the selected entries are not a random sample. All included entries were downloaded again and re-run with the current version.

## Results

### Benchmarks with known domains

RESUPER counted the domains correctly in all 13 synthetic ensembles (Jaccard 1.000), as did CYRANGE (0.998); the reference mode missed every three-body case (7/13). On the 134 curated real ensembles, RESUPER gave the same number of domains as CYRANGE for 130 and selected slightly fewer residues at a lower RMSD (coverage 0.818 vs 0.828; RMSD 0.544 vs 0.565 Å). On the frozen test sets, RESUPER counted the domains correctly in 95/96 (set 3) and 45/48 (set R) ensembles, against 76/96 and 39/48 for CYRANGE. It mixed two rigid bodies in one range in 0 and 4 ensembles, against 17 and 9, and it detected more hinges (Table 2, Figure 2). All criteria fixed in advance were met. On the multi-body set, RESUPER was correct in 155 of the 160 multi-domain cases and CYRANGE in 116; CYRANGE tends to combine several bodies into one range. A mixed range cannot be seen in a program’s own report. On set 3, CYRANGE counted the domains correctly in 76 of 96 ensembles (Jaccard 0.901), but 17 ensembles contained a range with residues from two separate rigid bodies, and nothing in the report distinguishes these 17 from the other 79.

**Figure 2.**
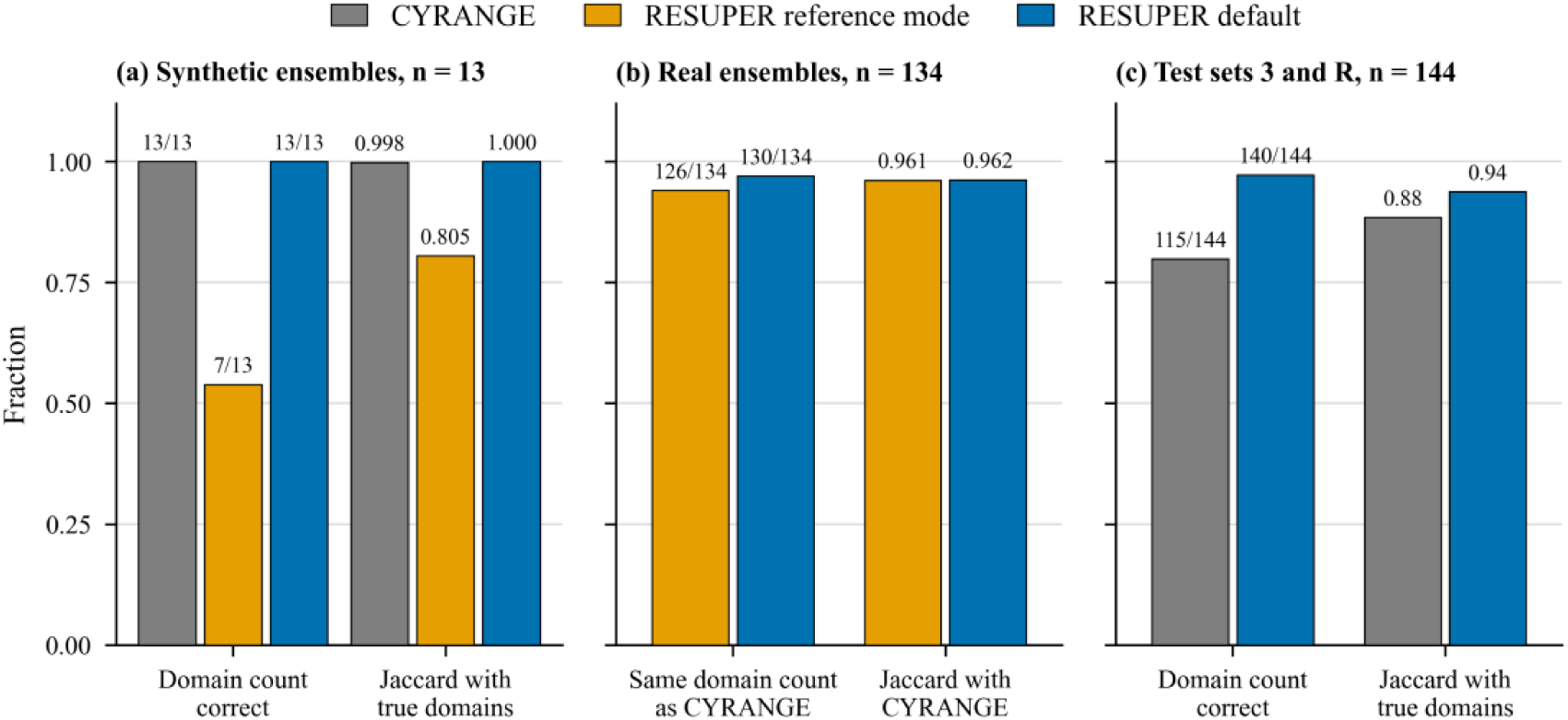
Benchmarks with known domains. (a) Ground truth on 13 synthetic ensembles. (b) Agreement with CYRANGE on 134 real NMR ensembles. (c) Ground truth on the frozen test sets 3 and R (144 ensembles). RESUPER is the default method; the reference mode is the CYRANGE-comparison mode.

**Table 2.** Benchmarks with known domains: domains counted correctly, with mixed domains or other measures in parentheses. Sets 1–2 were development data; sets 3 and R were run after the method was frozen.

| Benchmark | CYRANGE | Reference mode | RESUPER |
|---|---|---|---|
| Synthetic (13): counts (Jaccard) | 13/13 (0.998) | 7/13 (0.805) | 13/13 (1.000) |
| Curated real ensembles (134): same count as CYRANGE (RMSD) | – (0.565 Å) | 126 (0.578 Å) | 130 (0.544 Å) |
| Held-out set 1 (96): counts (mixed) | 74 (18) | 66 (24) | 96 (0) |
| Held-out set 2 (96): counts (mixed) | 76 (17) | 68 (21) | 94 (0) |
| Test set 3 (96): counts (mixed); hinges /56 | 76 (17); 39 | 68 (22); 33 | 95 (0); 43 |
| Test set R (48): counts (mixed); hinges /84 | 39 (9); 33 | – | 45 (4); 38 |
| Multi-body, 160 multi-domain cases of 788 ensembles: counts (mixed) | 116 (42) | 91 (63) | 155 (1) |
| Perturbation: ensembles changing count | 8 | 9 | 6 |

Under perturbation, the domain count changed for 6 ensembles with RESUPER, 8 with CYRANGE and 9 with the reference mode. Only RESUPER returned identical ranges for every ensemble when the model order was reversed, and CYRANGE failed to produce a report for one ensemble (2JQF) after coordinate noise was added. The domain count also changed little with the parameters: for τ = 0.06–0.20 Å it was unchanged for 128–131 of the 134 real ensembles, average linkage on the square root of the variance matrix gave the same decomposition as Ward linkage for all but one, and with any of four torsion sets the count stayed within 130–131.

### Comparison with CYRANGE: random samples

All 222 tested entries had a wwPDB CYRANGE result. RESUPER and CYRANGE gave identical domains for 136 entries (61%) and the same number of domains for 196 (88%). Agreement was highest for two-domain proteins (identical for 40 of 56, group A) and lowest for helical peptides (16 of 55, group E) and for proteins with three or more annotated domains (6 of 15, group B). The single-domain controls were in between (44 of 56, group C). None of the four hypotheses was significant, either on the confirmation sample of 162 entries or on all 222 entries (Table 3). CYRANGE recovered slightly more annotated domains: 38 against 32 in the confirmation sample (better in 4 proteins, RESUPER in none; p = 0.125) and 66 against 63 over all 222 (better in 6 proteins, RESUPER in 4; p = 0.678). Of the domains found by only one program, the rigid-domain check was passed by 7 of the 16 RESUPER-only domains and 18 of the 29 CYRANGE-only domains over all 222 (5 of 8 and 11 of 16 in the confirmation sample). Against the wwPDB CYRANGE results, the CYRANGE-only domains were therefore more often supported. The reference mode gave the same domains as CYRANGE for 158 of 222 entries, more often than the default method, but it over-split the single-domain controls (64 domains, against 57 for both CYRANGE and RESUPER) and merged domains in group B (30 domains, against 40 for CYRANGE). In the confirmation sample of 162 entries, the reference mode matched CYRANGE exactly in 117 entries and differed substantially in 34.

**Table 3.**
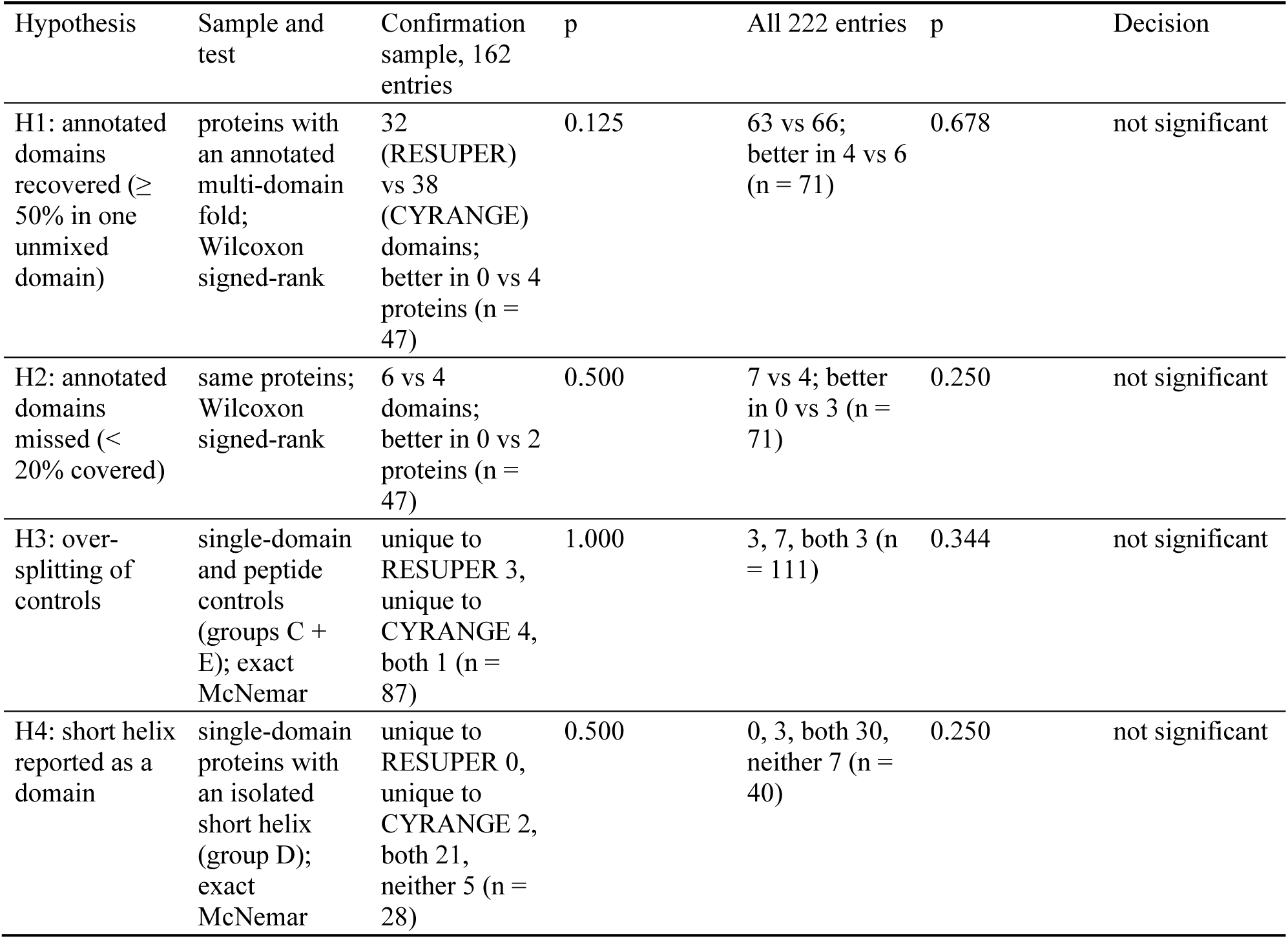
The four prespecified hypotheses run against the wwPDB’s CYRANGE domains. The hypotheses were specified with the reference mode as the comparator; applied to real CYRANGE the same rules are post hoc. All tests are two-sided at α = 0.05, as specified; no correction for the four comparisons was specified, and no decision would change under one.

| Hypothesis | Sample and test | Confirmation sample, 162 entries | p | All 222 entries | p | Decision |
|---|---|---|---|---|---|---|
| H1: annotated domains recovered (≥ 50% in one unmixed domain) | proteins with an annotated multi-domain fold; Wilcoxon signed-rank | 32 (RESUPER) vs 38 (CYRANGE) domains; better in 0 vs 4 proteins (n = 47) | 0.125 | 63 vs 66; better in 4 vs 6 (n = 71) | 0.678 | not significant |
| H2: annotated domains missed (< 20% covered) | same proteins; Wilcoxon signed-rank | 6 vs 4 domains; better in 0 vs 2 proteins (n = 47) | 0.500 | 7 vs 4; better in 0 vs 3 (n = 71) | 0.250 | not significant |
| H3: over-splitting of controls | single-domain and peptide controls (groups C + E); exact McNemar | unique to RESUPER 3, unique to CYRANGE 4, both 1 (n = 87) | 1.000 | 3, 7, both 3 (n = 111) | 0.344 | not significant |
| H4: short helix reported as a domain | single-domain proteins with an isolated short helix (group D); exact McNemar | unique to RESUPER 0, unique to CYRANGE 2, both 21, neither 5 (n = 28) | 0.500 | 0, 3, both 30, neither 7 (n = 40) | 0.250 | not significant |

### Illustrative cases

Of the 9,716 remaining pool entries, 8,709 were compared with CYRANGE; 693 were excluded by the screening rules, 303 had no CYRANGE result and 11 had no PDB-format file. The two programs differed substantially for 862 entries. The evidence rules were applied to 908 entries (these 862 and the 46 entries that differed in the two random samples) and supported RESUPER’s assignment in 164 of them, through annotation, a CYRANGE domain that is not internally rigid, or a RESUPER-only domain that is both rigid and moving. Additional large domains from RESUPER were rare. In only 17 of the 908 entries did RESUPER report more domains of ≥ 20 residues than CYRANGE; 7 of these met the category-A criteria, and in the other 10 the same evidence favored CYRANGE. One of the 7 was dropped because its own range RMSD was 1.87 Å, above the rigidity limit of the category, leaving 6. Of 42 category-B candidates, 32 were not included because the original publication did not describe the region explicitly, leaving 10. Table 4 lists these 16 entries. In category A, RESUPER separates domains that are described in the original publication and that CYRANGE does not report separately. In category B, the two programs assign the same domains, but CYRANGE also reports a region that the original publication describes as disordered or flexible. All 16 entries were re-run from freshly downloaded files with the current version, and every domain and RMSD was reproduced. These entries show where the two programs differ in RESUPER’s favor; the random samples above show how often this happens.

**Table 4.** Illustrative entries where independent evidence supports RESUPER’s decomposition. Category A: RESUPER reports more domains of ≥ 20 residues than CYRANGE, all internally rigid (≤ 1.0 Å), none splitting a single HELIX record, and the original publication does not contradict the division; CYRANGE does not report these domains separately. Category B: the two programs agree on every domain RESUPER reports (Jaccard ≥ 0.9) while CYRANGE additionally reports a region of ≥ 8 residues with range RMSD > 1.0 Å that the original publication explicitly describes as disordered or flexible. Selected for this property, so not a random sample. Ranges are given as first–last residue with the range RMSD, and multiple ranges of one domain are joined by +.

| Cat | PDB | Protein | Models, residues | RESUPER domains (range RMSD) | CYRANGE domains (range RMSD) | Supporting evidence; original publication |
|---|---|---|---|---|---|---|
| A | 2BW2 | Bypass of forespore c | 25, 140 | 8–61 (0.36 Å) + 64–140 (0.56 Å) | 7–140 (1.13 Å) | ECOD: 2 vs 0 annotated domains recovered; CYRANGE range non-rigid; paper<sup>23</sup>: two domains |
| A | 1BBO | Human enhancer-binding protein mbp-1 | 60, 56 | 2–10, 12–25 (0.19 Å) + 27–54 (0.26 Å) | 2–10, 12–55 (0.83 Å) | CATH: 2 vs 0; paper<sup>24</sup>: RMSD reported per zinc finger |
| A | 2EW9 | Copper-transporting ATPase 2 | 31, 149 | 4–13, 18–71 (0.42 Å) + 81–146 (0.46 Å) | 4–72, 80–149 (0.95 Å) | CATH: 2 vs 0; paper<sup>25</sup>: two ferredoxin folds, but they move as a unit (partial support) |
| A | 5MS9 | Fibrillin-1 | 20, 177 | 121–181 (0.80 Å) + 183–196, 200–287 (0.83 Å) | 140–287 (1.61 Å) | ECOD: 1 vs 0; CYRANGE range non-rigid; paper<sup>26</sup>: domains joined by a flexible pivot |
| A | 6G81 | Hydrogenase maturation factor HypA | 20, 117 | 2–69, 105–116 (0.32 Å) + 70–104 (0.38 Å) | 1–117 (0.73 Å) | RESUPER-only domain rigid and moving; paper<sup>27</sup>: Ni- and Zn-binding domains that reorient |
| A | 2V93 | Maltose-binding periplasmic protein | 50, 366 | 5–109, 262–319 (0.10 Å) + 114–253, 332–370 (0.08 Å) | 81–87, 103–105, 266–268, 311–313 (0.00 Å) | RESUPER-only domain rigid and moving; paper<sup>28</sup>: two domains, open/closed (PRE-refined ensemble) |
| B | 9FLG | Large delta antigen | 20, 140 | 99–134 (0.41 Å) | 60–75 (2.52 Å) + 99–134 (0.41 Å) + 162–179 (3.07 Å) | Extra CYRANGE domains 2.52, 3.07 Å; paper<sup>29</sup>: disordered regions flank the helix-loop-helix (96–142) |
| B | 1HN6 | Apical membrane antigen 1 | 20, 110 | 441–450, 481–509 (0.79 Å) | 441–450, 481–509 (0.79 Å) + 513–525 (2.24 Å) + 526–534 (1.31 Å) | Extra CYRANGE domains 2.24, 1.31 Å; paper<sup>30</sup>: N- and C-terminal regions unstructured |
| B | 2MCD | Murine norovirus 1 | 20, 98 | 62–112 (0.21 Å) | 11–11, 28–44 (2.22 Å) + 61–114 (0.22 Å) | Extra CYRANGE domain 2.22 Å; paper<sup>31</sup>: N-terminal PEST-like region unstructured |
| B | 7YWR | ORF1a polyprotein (Nsp8 N-terminal region) | 20, 84 | 4–55 (0.43 Å) | 4–54 (0.43 Å) + 55–77 (2.16 Å) | Extra CYRANGE domain 2.16 Å; paper<sup>32</sup>: helix fades after residue 50; 74–84 disordered |
| B | 1K9C | Calreticulin | 20, 74 | 214–258 (0.89 Å) | 195–206 (1.67 Å) + 214–261 (0.98 Å) | Extra CYRANGE domain 1.67 Å; paper<sup>33</sup>: residues 189–213 disordered |
| B | 2K8P | Sclerostin | 36, 189 | 57–82, 110–125, 130–144 (0.48 Å) | 8–21 (1.57 Å) + 29–39 (1.18 Å) + 57–82, 110–125, 130–144 (0.48 Å) + 158–173 (1.33 Å) | Extra CYRANGE domains 1.57, 1.18, 1.33 Å; paper<sup>34</sup>: N- and C-terminal arms unstructured, highly flexible |
| B | 2PXG | Outer membrane protein | 20, 118 | 18–56, 70–88 (0.93 Å) | 18–56, 70–88 (0.93 Å) + 105–112 (1.56 Å) | Extra CYRANGE domain 1.56 Å; paper<sup>35</sup>: N- and C-termini highly flexible |
| B | 2LWR | SD01679p | 19, 68 | 431–481 (0.82 Å) | 418–427 (1.52 Å) + 431–482 (0.83 Å) | Extra CYRANGE domain 1.52 Å; paper<sup>36</sup>: N terminus disordered (A417–I429) |
| B | 2KTV | Eukaryotic peptide chain release factor subunit 1 | 24, 164 | 278–334, 342–416 (0.54 Å) | 278–334, 342–416 (0.54 Å) + 421–440 (1.44 Å) | Extra CYRANGE domain 1.44 Å; paper<sup>37</sup>: C-terminal tail 414–437 flexible (<sup>15</sup>N relaxation) |
| B | 5NR5 | MatA protein | 30, 113 | 35–77 (0.27 Å) | 1–12 (1.05 Å) + 35–77 (0.27 Å) + 95–103 (1.25 Å) | Extra CYRANGE domains 1.05, 1.25 Å; paper<sup>38</sup>: N- and C-terminal tails largely disordered |

### Multiconformer detection: 7SA5

Apo-Pin1 (PDB 7SA5) was deposited as a two-state ensemble of 20 conformers^39^, with the conformers of the two states interleaved in model order. RESUPER and CYRANGE assign the same two domains (WW, residues 5–39; PPIase, 53–163), so the domain ranges do not reveal the states. The detector reports two states of equal size (silhouette 0.745), models 2, 4–5, 8–9, 12–13, 16, 18, 20 and models 1, 3, 6–7, 10–11, 14–15, 17, 19 (Figure 1a). The Pin1 complexes 7SUQ^40^, 7SUR^40^ and 8SG2^41^ were also reported as two states (silhouette 0.744, 0.667 and 0.622), and single-state controls as one state. Because the 0.60 threshold was calibrated on these Pin1 ensembles, the results illustrate the detector but do not validate it independently. The two states differ in the arrangement of the two domains, not in their internal structure. Measured in each conformer, the distance between the Cα centroids of the two reported ranges is 21.7 ± 0.5 Å in one state and 24.5 ± 3.4 Å in the other (23.1 ± 2.8 Å over the whole ensemble). For 7SUQ the values are 22.1 ± 0.5 and 27.5 ± 0.9 Å, and for 7SUR 21.8 ± 0.5 and 27.1 ± 1.3 Å. Superimposing each state on the ranges of the whole ensemble leaves the range RMSD nearly unchanged (residues 5–39: 0.92 Å for the whole ensemble, 0.94 and 0.87 Å for the two states; residues 53–163: 0.87, 0.84 and 0.78 Å), so each domain has a similar internal structure in both states.

Analyzed state by state (`--per-state`), the two states give different ranges (Figure 1a, c, d). In state 1, where the domain centroids stay 21.7 ± 0.5 Å apart, RESUPER reports a single range that covers both domains (residues 4–38 and 52–163, RMSD 1.21 Å). In state 2 it reports the two domains separately (5–39, 0.87 Å; 53–163, 0.78 Å). The single range indicates a nearly fixed interdomain arrangement in state 1, not a strictly rigid one: its RMSD is above the 1.0 Å used for the rigid-domain check, and the angle between the principal axes of the two domains still varies (46.5 ± 7.1°, against 62.8 ± 16.2° in state 2).

7SUQ and 7SUR show the same pattern. In both, the state with the shorter and less variable domain separation also has the smaller spread of the interdomain angle (7SUQ: ± 6.2° against ± 15.5°; 7SUR: ± 6.4° against ± 9.5°). RESUPER reports a single range in that state (7SUQ: 10–37 and 52–163, 1.30 Å; 7SUR: 10–34 and 51–163, 0.99 Å) and two ranges in the other (7SUQ: 6–38, 0.92 Å, and 52–163, 1.05 Å; 7SUR: 5–39, 0.89 Å, and 52–163, 0.77 Å).

## Discussion

### Domain decisions

Each domain decision in RESUPER is based on a quantity in ångströms or a ratio: the split score of Eq. (3), the block structure of the distance-variance matrix, or a minimum domain size. A user can recompute these values for the structure at hand, which is not possible with a tuned constant. On the benchmarks with known domains, RESUPER differed from CYRANGE mainly in ensembles whose bodies move relative to each other: it was correct in 155 of 160 multi-domain cases against 116, and it mixed rigid bodies in 0 and 4 frozen-test ensembles against 17 and 9.

### Agreement with CYRANGE on deposited entries

On randomly selected NMR entries, the two programs mostly agreed, and none of the four prespecified hypotheses, applied post hoc to CYRANGE, reached significance. CYRANGE recovered slightly more annotated domains, and more of its unique domains passed the rigid-domain check. The main systematic difference was for helical peptides, where the programs gave identical domains for only 16 of 55 entries. In these entries, RESUPER often reports two ordered segments within what the deposited entry annotates as one helix. A continuous helix does not necessarily constitute a single rigidly superposable region across an NMR ensemble. In all 21 of the 908 differing entries in which a RESUPER split lies inside one deposited HELIX record, the relative orientation of the two segments varied across the conformers more than that of the two halves of the longer segment. Whether such motion within a helix should give separate superposition ranges depends on the use, and secondary-structure annotation cannot decide it. The selection rules for Table 4 therefore exclude these entries: a split inside a single deposited HELIX record disqualifies an entry. RESUPER’s advantage on the synthetic and simulated benchmarks therefore applies to cases where a body was moved deliberately and its boundaries are known. On routine experimental ensembles the two programs largely agree, and where they differ, fold annotation is too coarse to decide which is right. The entries in Table 4 show specific cases where independent evidence (fold annotation, domain rigidity, relative motion, or a statement in the original publication) supports one decomposition over the other. They were selected for this property and are not a random sample.

### Conformational states

The state detector answers a different question from the domain ranges. In 7SA5, each domain has a similar internal structure in both states, but the domain centroids are on average 2.8 Å further apart in state 2, and only in state 1 do the two domains share one superposition range, consistent with a nearly fixed interdomain arrangement. A superposition range computed over the whole ensemble reports none of this. The detector is simple: Ward partitions of a three-component distance descriptor, scored by the silhouette coefficient with a threshold of 0.60. The threshold was calibrated on the same Pin1 ensembles shown here, so these results are a demonstration rather than an independent validation. Every state must contain at least three conformers, so a state populated by fewer conformers is not reported, and a 20-conformer ensemble can be divided into at most six states. A state describes a pattern of Cα–Cα distances, not exchange rates or thermodynamic populations.

### Limitations

The thresholds τ, *g* and *d* have a geometric meaning but were chosen on development data; the frozen test sets show that they transfer, not that they are optimal. For some helical peptides, RESUPER identifies two separately superposable regions within a continuous helix, whereas CYRANGE reports a single domain. Such divisions may reflect relative motion between locally ordered segments rather than over-splitting, but independent ground truth is unavailable to determine which decomposition is more appropriate. Across the 222 randomly selected NMR entries, the programs differed in domain count for 26 entries and substantially in domain assignment for 46. The benchmarks with known domains rely on modeled hinge and linker motions, and the mix of scenarios, although fixed before evaluation, was designed by us. Ensembles with independently established domain motions, such as the Pin1 series, therefore provide a stronger test of both stages.

## Conclusion

RESUPER selects superposition ranges and counts conformational states in protein ensembles. It keeps the two-stage framework of CYRANGE and replaces the tuned steps of domain selection with geometric tests. On test sets fixed in advance, it counted the domains correctly in 95/96 and 45/48 ensembles (CYRANGE: 76/96 and 39/48) and mixed rigid bodies in 0 and 4 ensembles (CYRANGE: 17 and 9). On 222 randomly selected NMR entries, the two programs gave the same domain count for 88% of entries, so the difference lies mainly in ensembles whose bodies move independently. The state detector found the deposited two-state ensembles of Pin1, whose states differ in the relative position of the WW and PPIase domains; analyzed state by state, the two domains share one superposition range in one state and have separate ranges in the other. RESUPER is available in POKY with a graphical interface, CYRANGE-compatible command-line parameters, mmCIF support, per-state output, and range strings for Xplor-NIH, CYANA and PyMOL. It will be used across the Integrative NMR platform of POKY and, in the AUDANA and AUDASA structure calculations of PONDEROSA-C/S, to check the ordered regions of the ensemble at every cycle.

## Acknowledgements

This work was supported by the National Science Foundation [DBI-2413041 and DBI-2051595 to W.L.] and the University of Colorado Denver.

## Software availability

RESUPER is distributed with POKY, which can be downloaded from https://poky.clas.ucdenver.edu; in POKY it is opened from the RESUPER dialog (Figure 1; two-letter code *yS*). The source code, the tests, the benchmark scripts, the synthetic-ensemble generators and the frozen test-set designs are available at https://github.com/pokynmr/RESUPER_deploy.

